# Knockout of αβ but not γδ T cells in chickens is associated with high cytotoxicity and deficiency of regulatory and helper T cells

**DOI:** 10.1101/2023.03.10.531286

**Authors:** Theresa von Heyl, Romina Klinger, Dorothea Aumann, Christian Zenner, Mohanned Alhussien, Antonina Schlickenrieder, Kamila Lengyel, Hanna-Kaisa Vikkula, Teresa Mittermair, Hicham Sid, Benjamin Schusser

## Abstract

The availability of genetically modified mice has facilitated the study of mammalian T cells. No model has yet been developed to study these cells in chicken, an important livestock species with a high availability of γδ T cells. To investigate the role of γδ and αβ T cell populations in birds, we generated chickens lacking these T cell populations. This was achieved by genomic deletion of the constant region of the T cell receptor γ or αβ chain, leading to a complete loss of either γδ or αβ T cells. Our results show that a deletion of αβ T cells but not γδ T cells resulted in a severe phenotype in knockout chickens. The αβ T cell knockout chickens exhibited granulomas associated with inflammation of the spleen and the proventriculus. Immunophenotyping of αβ T cell knockout chickens revealed a significant increase in monocytes and the absence of CD4^+^ T cells and FoxP3^+^ regulatory T cells compared to wild type chickens. In addition, we observed a significant decrease in immunoglobulins, B lymphocytes, and changes in the bursa morphology. Our data reveal the consequences of T cell knockouts in chickens and provide new insights into their function in vertebrates.

**Significance statement:** The lack of genetically modified chickens has severely limited research in avian immunology compared to other animal models. Here, we report the generation of two T cell knockout chicken lines that will contribute significantly to the understanding of T cell biology as a very important research model as well as an important livestock species. The generated animals reveal the function of different T cell populations in chickens and will help to better understand the role of these cells during the interaction with various pathogens in birds.

## Introduction

T lymphocytes can recognize a variety of peptides to facilitate the humoral and cytotoxic immune responses and are greatly involved in variable processes including inflammatory and autoimmune diseases. They are characterized by their heterodimeric T cell receptor (TCR), which consists of a constant and a variable region with either αβ or γδ chains. Therefore T lymphocytes are divided into two major subgroups, the αβ and γδ T cells (1). The role of these cells has been largely studied in humans and mammalian animal models including mice, which harbor a comparatively low percentage of peripheral blood γδ T cells (2). In contrast, chickens harbor a high percentage of γδ T cells, which makes them an intriguing research model.

γδ T cells in chickens are known for their cytotoxic activity (3), and high availability in the peripheral epithelial tissue (5). The function of αβ T cells is defined by the two co-receptors CD4 and CD8. The CD4^+^ αβ T cells are activated through antigens bound to major histocompatibility complex (MHC) II molecules, leading to a humoral immune response against pathogens (6). Regulatory T cells (T_regs_), typically expressing CD4^+^CD25^+^ and the transcription factor FoxP3^+^ (7), play an important role in maintaining immune tolerance (8). CD8^+^ αβ T cells recognize peptides presented by MHC I molecules (9) and show cytotoxic activity (6). Similar to other species, chicken CD8^+^ T cells have two isoforms, composed of a CD8αα homodimer or a CD8αβ heterodimer. They can further be divided into CD8^+high^, CD8^+dim,^ and CD8^-neg^ populations, which were shown to respond differently to pathogens such as salmonella (4).

Due to the unavailability of genetically modified avian models, early research was based on thymectomy combined with monoclonal antibody injections to investigate chicken T cell biology, which can be associated with significant alterations of the immune system (10). Using this method it was not possible to define T cell functions since the frequency of epithelium-associated γδ T cells (TCR1) as well as Vβ1^+^ αβ T cells (TCR2) cells were not affected (10). More recently, the generation of recombination activating gene 1 (RAG1) knockout (KO) chickens led to the loss of both B and T cell populations (11) which helped in understanding the role of this gene in chicken lymphocyte development, but it did not allow a separate study of distinct immune cell populations.

The establishment of a reverse genetic system in chickens allowed the generation of a KO of the B cell receptor (BCR) using the genetic modification of primordial germ cells (PGCs) (12, 13). Not only the use of PGCs but also the availability of the Clustered Regularly Interspaced Short Palindromic Repeats (CRISPR)/Cas9 technology has facilitated the generation of genetically edited chickens and simplified the gene editing process (14). To define specific T cell populations’ functions, such as αβ T cells and γδ T cells, genetically modified chicken lines lacking these populations are pivotal.

In this study, we successfully generated both TCR Cγ^-/-^ and TCR Cβ^-/-^ chickens, lacking γδ or αβ T cells, respectively. TCR Cβ^-/-^ chickens exhibited a severe phenotype that resulted in inflammation in the spleen, proventriculus, and skin associated with structural deterioration of the thymus and bursa of Fabricius as early as 14 days after hatch, while no adverse phenotype was observed in TCR Cγ^-/-^ chickens. The observed phenotype in TCR Cβ^-/-^ indicates an autoimmune condition caused by the loss of CD4^+^ helper T cells and FoxP3^+^ regulatory T cells.

## Results

### Deletion of the T Cell receptor β and γ constant chains

Chickens lacking either γδ or αβ T cells were generated by targeting the γ or respectively β constant region of the corresponding TCR. CRISPR/Cas9 mediated homology-directed repair (HDR) was used in chicken PGCs to replace the constant region by a selectable marker cassette (Fig. 1*A,B*). Precise targeting was confirmed by knockout allele-specific PCR and correctly targeted clonal PGC lines were used to generate TCR Cγ^+/-^ (germline transmission rate = 5.3%) and TCR Cβ^+/-^ (germline transmission rate = 1.5%) animals (Fig. S1*A*). To remove the selectable marker cassette, PGCs were re-derived from these lines and electroporated with Cre recombinase to remove the selectable marker cassette. eGFP^neg^ PGCs were sorted by FACS and injected into embryos to finally generate TCR Cγ^+/-^ (germline transmission rate = 9%) and TCR Cβ^+/-^ (germline transmission rate = 5,9%) chickens that do not express eGFP (Fig. 1*A,B*; Fig. S1*A*). Knockout and wild type allele-specific PCRs were established to detect TCR Cγ^-/-^ and TCR Cβ^-/-^ chickens (Fig. 1*C,D*).

**Fig. 1:**
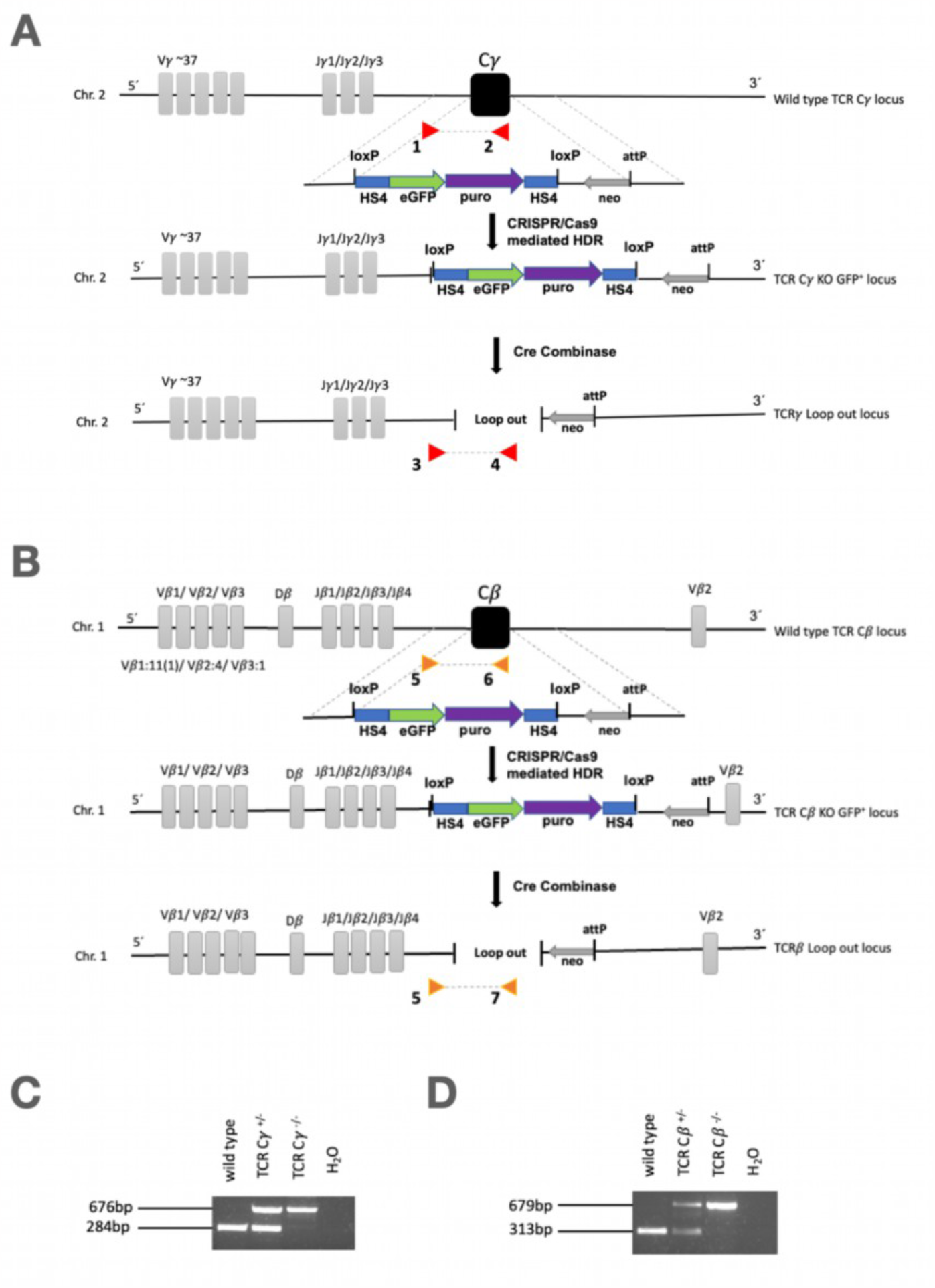
Generation of the TCR C*β*^-/-^ and TCR C*γ*^-/-^ knockout animals. (A) The establishment of a γδ T cell deficient chicken line was achieved by targeting the constant region of the TCR chain on chromosome 2. The targeting vector contained a selectable marker cassette (eGFP + puromycin resistance gene) to allow the selection of successfully targeted cells. Cre-mediated recombination resulted in the removal of the selectable markers. (B) The establishment of an αβ T cell deficient chicken line was achieved by targeting the constant region of the TCR β chain on chromosome 1. The same targeting strategy as shown for the TCR chain was used. (C) Genotyping of TCR Cγ^-/-^ chicken by multiplex PCR. Wild type allele-specific primers (#1 and #2) with an amplicon length of 284bp and TCR C knockout-specific primers (#3 and #4) with an amplicon length of 676bp were used. (D) Genotyping of TCR C*β*^-/-^ chicken by multiplex PCR. Wild type allele-specific primers (#5 and #6) with an amplicon length of 313bp and TCR Cβ KO-specific primers (#5 and #7) with an amplicon length of 679bp were used.

### TCR Cγ^-/-^ and TCR Cβ^-/-^ chickens lack either γδ or αβ T cells

The immunophenotype of the generated T cell knockout chicken was analyzed via flow cytometry. TCR Cγ^-/-^ chickens showed a complete depletion of γδ T cells and TCR Cβ^-/-^ chickens a complete depletion of αβ T cells. Simultaneous staining for the pan T cell antigen CD3 confirmed the successful knockout and revealed that no uncharacterized T cell population exists in chickens (Fig. 2*A,B*). While TCR Cγ^-/-^ chickens are phenotypically similar to wild type animals, TCR Cβ^-/-^ chickens develop a severe phenotype as early as two weeks post hatch. Macroscopic lesions like epithelial granulomas at the comb, beak, and legs associated with inflammation in the mucosal surface of the proventriculus (Fig. S1*D*) and the spleen (Fig. 4*A*) were seen. To investigate whether the phenotype was caused to environmental pathogens or autoimmune factors, biopsy samples from the spleen of two-week-old TCR Cβ^-/-^ animals and wild type animals were cultivated for bacterial detection. No bacterial growth was observed, confirming that the phenotype was not related by environmental infection (Data not shown). Neither TCR Cγ^-/-^ nor TCR Cβ^-/-^ chickens showed impaired weights compared to their wild type siblings (Fig. S1*B,C*).

**Fig. 2:**
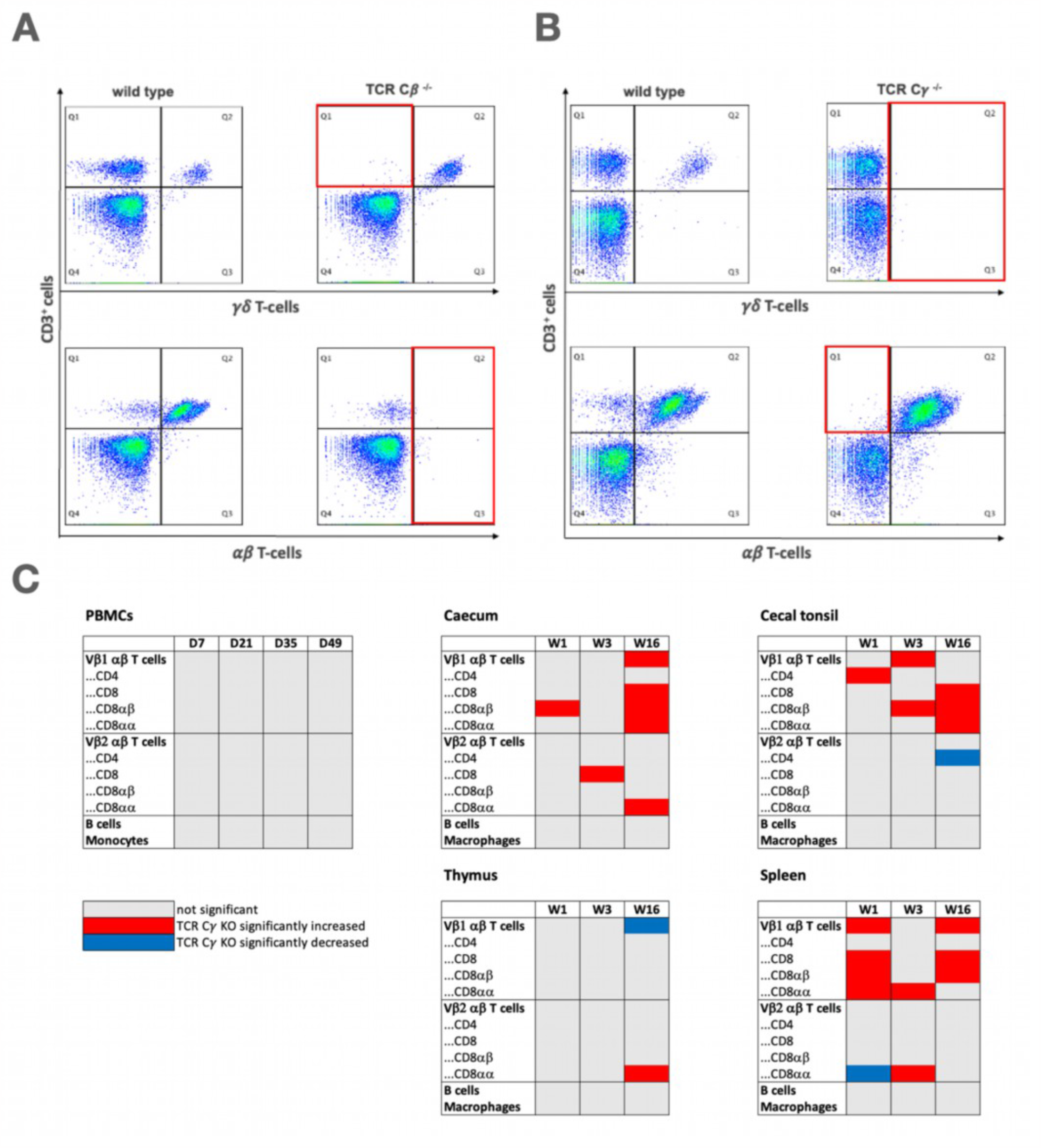
(A) One representative dot plot of wild type against TCR C*β*^-/-^ (day 14; n=10) and (B) wild type against TCR Cγ^-/-^ knockout chickens (day 24; n=3) is shown. PBMCs were stained for the T cell marker CD3 and γδ T cells (TCR1) or αβ T cells (TCR2 + TCR3). (C) Analysis of TCR2 subsets, TCR3 subsets, B cells, and macrophages at day (D) 7, 21, 35, and 49 from PBMCs and mononuclear cells isolated from thymus, spleen, caecum and cecal tonsils at week (W) 1, 3, and 16 of TCR Cγ^-/-^ knockout chickens (n=4). p<0.05

### The absence of γδ T cells is compensated through CD8^+^ αβ T cells in the gut

Analysis of peripheral blood mononuclear cells (PBMCs) in TCR Cγ^-/-^ chickens showed, that the loss of γδ T cells does not influence other blood circulating T cell subpopulations, B cells, or monocytes between seven- and 49 days post-hatch. However, a significantly higher number of Vβ1^+^ αβ T cells was found in the spleen after one and 16 weeks (*p<0*.*05*). Within these, the CD8^+^ T cells in both isotypes CD8αβ^+^ and CD8αα^+^ were increased. In the caecum, significantly higher number of CD8αβ^+^ Vβ2^+^ αβ T cells was found seven days post hatch, and a significantly higher number of CD8^+^ Vβ1^+^ αβ T cells was found after 16 weeks (*p<0*.*05*). In the cecal tonsils, significantly more CD4^+^ Vβ1^+^ αβ T cells were detected (*p<0*.*05*) after one week. However, after three weeks the CD8αβ^+^ Vβ1^+^ αβ T cells were increased (*p<0*.*05*), and after 16 weeks both isotypes CD8αα^+^ and CD8αβ^+^ from Vβ1^+^ αβ T cells were increased (*p<0*.*05*). In the thymus, the Vβ1^+^ αβ T cells were decreased (*p<0*.*05*), and the CD8αα^+^ Vβ2^+^ αβ T cells were increased (*p<0*.*05*) after 16 weeks (Fig. 2C). The intestinal integrity was examined by morphometric analysis of histological sections including the length of the tunica muscularis, crypts, and villi in the duodenum, jejunum, ileum, and caecum of TCR Cγ^-/-^ animals compared to wild type birds. No significant differences were found between both groups (*p>0*.*05*) (Fig. S6).

### TCR Cβ^-/-^ chickens show an increase in monocytes and a decrease in B cells

In the TCR Cβ^-/-^ chicken no compensatory effect of the γδ T cells was detected. 14 days after hatch significantly fewer B cells, CD4^+^, and CD8^+^ T cells were detected (*p<0*.*05*) along with significantly increased numbers of monocytes of TCR Cβ^-/-^ chicken (*p<0*.*05*) (Fig. 3*A*). The number of blood circulating γδ T cells was significantly lower in double positive CD8^+dim^ CD4^+^ cells, one week after hatch (p<0.05). On the other side, levels of both CD8^+high^ and CD8^+dim^ cells within the γδT cells were significantly higher after seven and 15 days post-hatch (Fig. 3*B*).

**Fig. 3:**
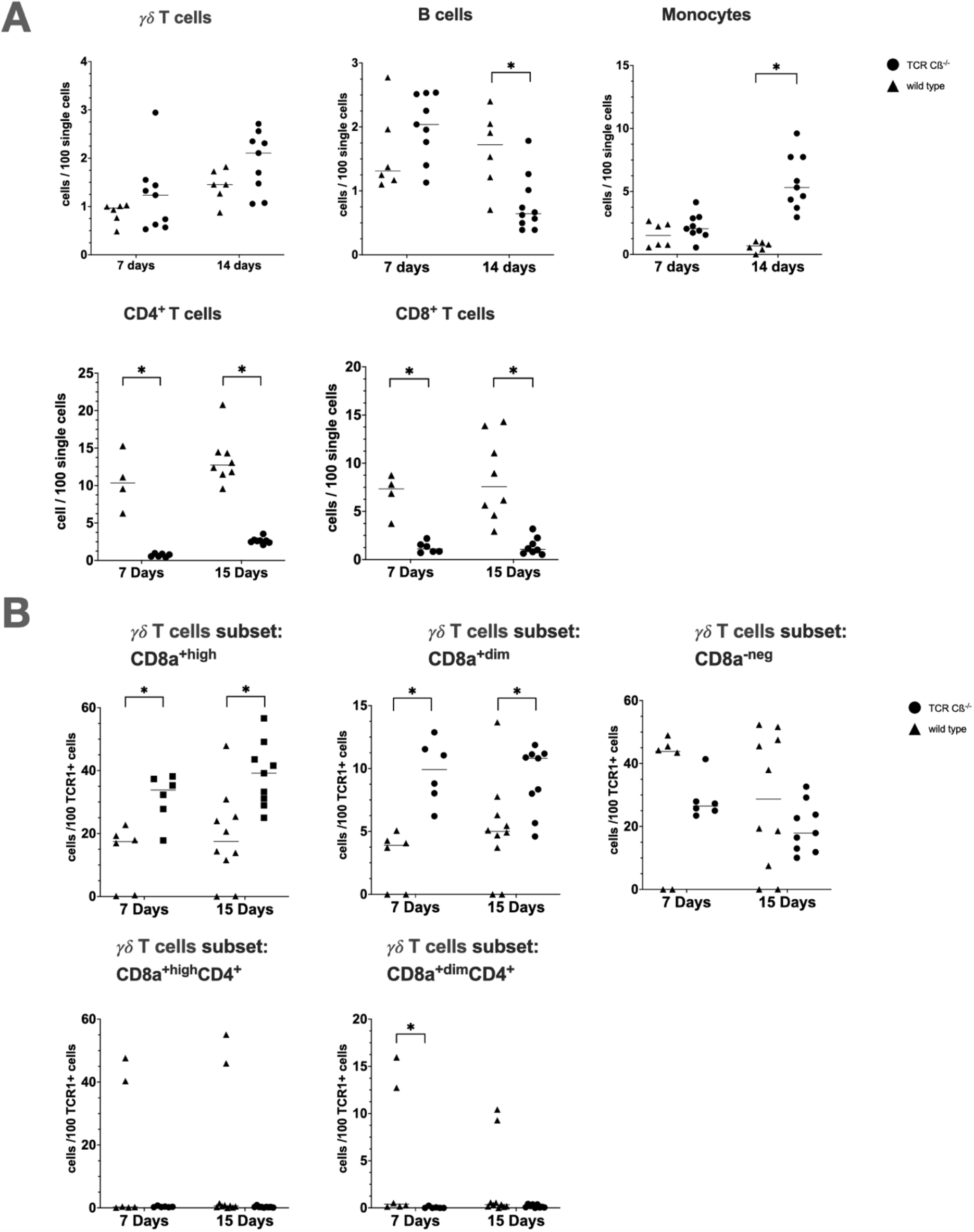
(A) Flow cytometric analysis of PBMCs of TCR C*β*^-/-^ (n≥6) and wild type (n≥4) chickens at seven and 14 days of age. (B) CD8^+^ cells within the γδ T cell (TCR1) subset were analyzed based on their level of CD8 expression and co-expression of CD4 from TCR C*β*^-/-^ (n≥6) and wild type (n≥4) chickens at seven and 15 days of age. Mean and standard deviation of n*≥* 4 are shown. * p<0.05

Additionally, immunoglobulin levels in the plasma of 14 day-old TCR Cβ^-/-^ chickens were decreased compared to wild type chickens while no differences between TCR Cγ^-/-^ and wild type chickens were observed (Fig. S2).

### TCR Cβ^-/-^ knockout impacts the development of lymphatic organs

The thymus in wild type chickens showed a clear separation in cortex and medulla. No such separation was found in TCR Cβ^-/-^ chickens (Fig. 4*A*). The spleen of TCR Cβ^-/-^ chickens showed inflammatory lesions (Fig. 4*A*). Surprisingly, a strong phenotype was observed in the bursa of Fabricius in TCR Cβ^-/-^ chickens, where B cell follicles appeared underdeveloped in comparison to wild type siblings (Fig. 4*A*). Immunohistochemistry of the epithelial granulomas 14 days after hatch showed infiltration of the surrounding tissue by macrophages and a central granulocyte accumulation, surrounded with B cells and γδ T cells (Fig. 4*B*). Splenic immunofluorescence staining of B cells, CD4^+^, and CD8^+^ T cells revealed the absence of germinal center formations and CD4^+^ T cells in TCR Cβ^-/-^ animals, which was not the case in wild type birds. In addition, TCR Cβ^-/-^ chickens exhibited a random splenic distribution of B cells and a few CD8^+^ T cells, while typical white pulp formation of B cells surrounded by CD8^+^ T cells was found in wild type animals (Fig. S3).

**Fig. 4:**
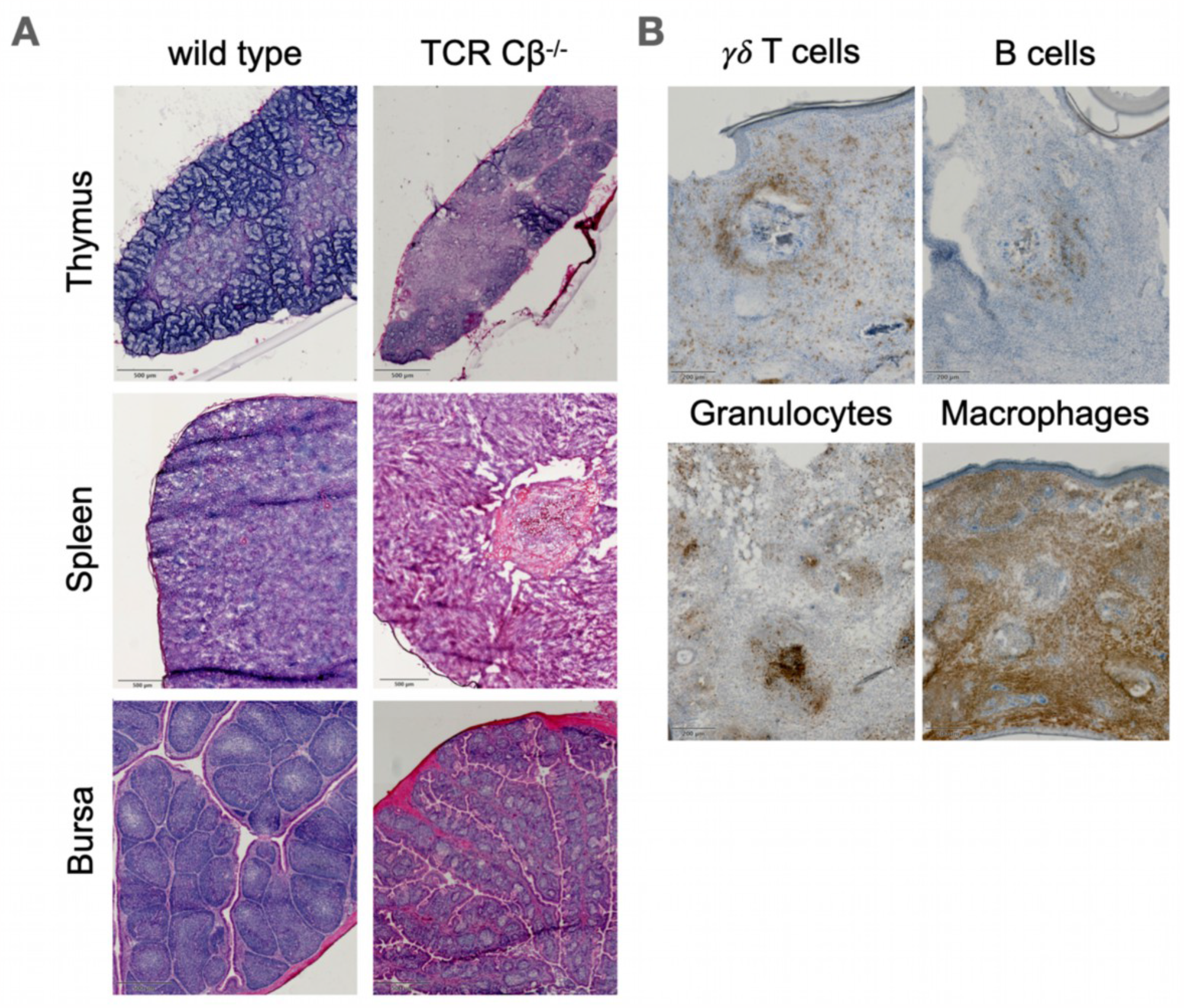
(A) Sections of thymus, spleen, and bursa of 14-day-old wild type and TCR C*β*^-/-^ chicken have been prepared and stained with H&E. (B) Skin granulomas of 14-day-old wild type and TCR C*β*^-/-^ chicken have been dissected and stained for γδ T cells (TCR1), B cells (BU1/AV20), granulocytes (GRL1) and macrophages (Kul01). Always one representative picture of at least three different animals per genotype is shown. Scale bar (A) = 500µm (B) = 200µm.

### FoxP3^+^ expression of T_reg_ cells is significantly lower in TCR Cβ^-/-^ knockout chicken

The expression of several immune-related genes was compared between wild type, TCR Cβ^-/-^ and TCR Cγ^-/-^ chickens, 14 days after hatch. Significantly lower TGFβ and IL-5 expression was found in the spleen (*p<0*.*05*), while significantly higher IL-6 levels were detected in the thymus of TCR Cβ^-/-^ animals (*p<0*.*05*). In addition, a significantly increased IL-1ß expression was detected in PBMCs of TCR Cβ^-/-^ animals (*p<0*.*05*). IL-22 expression was significantly lower in the spleen of TCR Cβ^-/-^ chickens, while it was significantly higher in the spleen of TCR Cγ^-/-^ chickens (*p<0*.*05*).

Following the significantly decreased CD4^+^ T cell population FoxP3 expression was significantly lower in the spleen, thymus, and PBMCs in TCR Cβ^-/-^ animals (*p<0*.*05*) (Fig. S4).

### The gut microbiome of TCR Cβ^-/-^ chickens shows a significantly different beta diversity

To analyze the influence of T cells in the chicken gut microbiome, 16S RNA sequencing of feces and caecum content was performed. Significant differences in the number of species were found in the feces of 14-day-old TCR Cβ^-/-^ chickens as well as in the caecum of 35-day-old TCR Cγ^-/-^ chickens (*p<0*.*05*). There were no differences in the Shannon effective counts (*p*>0.05). Analyzing the beta diversity of both fecal and cecal samples of TCR Cβ^-/-^ chickens demonstrated significantly different populations compared to wild type chickens (*p<0*.*05*). In the caecum content of 14-day-old TCR Cβ^-/-^ chickens, significantly fewer bacteria of the class Clostridia were found, while in the feces, significantly fewer Actinobacteria and significantly more Clostridia were detected at the class level (*p<0*.*05*) (Fig. S5).

## Discussion

T cells play a crucial role in regulating various physiological functions and providing defense against pathogens in humans and other mammals (1). However, the role of each subpopulation in chickens remains unclear due to the lack of investigative tools such as genetically engineered chickens in the past. This study was carried out to characterize the functions of different T cell subpopulations by generating chickens that lack γδ T cells (TCR Cγ^-/-^) and those that lack αβ T cells (TCR Cβ^-/-^). The successful generation and detailed characterization of these lines showed that the loss of γδ T cells in TCR Cγ^-/-^ chicken was asymptomatic, while the TCR Cβ^-/-^ chickens exhibited a severe phenotype.

It is widely believed that γδ T cells play a central role in mediating various functions of the immune system in chickens since they harbor a high number of γδ T cells with up to 50% in blood (5, 12). Surprisingly, the TCR Cγ^-/-^ chickens did not display major phenotypic differences compared to wild type chickens. In mice, a significant population of γδ T cells is present in the intestine and plays an important role in maintaining intestinal integrity (15). Contrary to expectations no differences were recorded in the morphometry of the intestine in the TCR Cγ^-/-^ compared to wild type chickens. It is well established that the majority of γδ T cells in chickens are CD8^+^ T cells (4). In TCR Cγ^-/-^ chickens the CD8^+^ αβ T cells population is significantly increased in the caecum, spleen, and cecal tonsils. Concluding that this CD8^+^ αβ T cell population compensates for the absence of the γδ T cells in the gut and therefore, maintains intestinal integrity in TCR Cγ^-/-^ chickens. This phenomenon is also supported by the findings of Sandrock et al. (2018), who reported that γδ T cell-deficient mice show mild phenotypes and that other lymphocytes take over the functions of γδ T cells in the absence of these cells (16).

The TCR Cβ^-/-^ chicken developed a pathological phenotype around two weeks of age. To determine if the observed phenotype is caused by a microbial infection, the spleens of 14-day-old TCR Cβ^-/-^ and wild type chickens were cultured to check for bacterial growth, but no bacteria were detected after a week of cultivation. The findings of this study indicate that the severe phenotype of the TCR Cβ^-/-^ is caused autoimmune due to an imbalance of humoral and cytotoxic responses and the loss of CD4^+^ T cells and T^reg^ cells. This phenotype was not in agreement with Cihak et. al. (1993) who reported depletion of T cell subsets after thymectomy and antibody injections without a pronounced phenotype. The method employed by Cihak et. al. was deemed inefficient, as the combination of TCR2 antibody treatment and thymectomy led to an increase of TCR3^+^ cells, thus demonstrating the inability to effectively eliminate T cells completely (10). Later research showed, that mice with TCR ^-/-^ KO also develop an inflammation of the stomach (17). Likewise, severe combined immunodeficiency (SCID) mice and RAG1 deficient chicken show similar phenotypes to the TCR Cβ^-/-^ chickens. Even though in SCID mice and Rag-1 deficient chickens it is suggested that the severe phenotype appears through the loss of both B- and T cells (11, 18), we showed in this study, that the phenotype in chickens appears only because of the loss of αβ T cells. Also, chickens lacking B cells (12) and TCR Cγ^-/-^ chickens do not show this phenotype, underlining that only the missing αβ T cells are leading to this severe phenotype.

The significant decrease in CD4^+^ T cells in the PBMCs of TCR Cβ^-/-^ chicken is one of the key findings to explain the severe pathology of this phenotype. CD4^+^ T cells secrete anti-inflammatory cytokines to coordinate the functions of other immune cells, primarily B cells, macrophages, and cytotoxic T cells (9, 19). This helps to prevent an overactive immune response that can result in autoimmune diseases (20).

Indeed, the TCR Cβ^-/-^ chickens showed a reduction in the expression of IL-4, and IL-5, a significant decrease of B cells and a concurrent reduction of IgY, IgM, and IgA compared to wild type chickens. These observations may be attributed to the absence of CD4^+^ T cells, as the secretion of IL-4 and IL-5 controls the activation and proliferation of B cells and therefore the production of immunoglobulins (16).

The lower IL-22 mRNA expression especially in spleen of TCR Cβ^-/-^ chickens compared to wild type chickens can be explained by the fact that CD4^+^ T cells are a major source of IL-22 in chickens (21). IL-22 plays a critical role in enhancing the innate immunity of tissues and facilitating repair and healing mechanisms during inflammation, which is essential for restoring tissue homeostasis and preventing autoimmune diseases (22). This could also contribute to the phenotype observed in TCR Cβ^-/-^ chickens.

Moreover, the significantly lower expression of FoxP3 and TGFβ in TCR Cβ^-/-^ chickens matches the absence of CD4^+^ T cells and also indicates the loss of T_reg_ cells. IL-10 and TGFβ secreted by T_reg_ cells have anti-inflammatory and immunosuppressive effects, inhibiting the activation and function of CD8^+^ T cells and monocytes, which are involved in the cytotoxic response (7). In the TCR Cβ^-/-^ chicken the regulation of cytotoxic activity of γδ T cells and macrophages is therefore disrupted.

In total fewer CD8^+^ T cells were found in the PBMCs of the TCR Cβ^-/-^ chickens. But within the TCR1 subsets, both CD8^+high^ and CD8^+dim^ T cells were significantly increased. Whereas after infection with *Salmonella enterica*, the CD8^+high^ TCR1^+^ population increases while the CD8^+dim^ TCR1^+^ subset decreases (4). In TCR Cβ^-/-^ chicken the majority of TCR1^+^ T cells are CD8^+^ (23), leading to a disbalance of the cytotoxic and humoral response and therefore higher levels of both CD8^+high^ and CD8^+dim^ T cells. Interestingly, even though it has been described, that double-positive CD8^+^CD4^+^ T cells are increasing in autoimmune diseases, for example, thyroiditis (24), in the TCR Cβ^-/-^ chickens fewer double-positive T cells were seen. It is still open to investigate whether this is related to the loss of CD4^+^ T cells in general. The increase in the monocyte population and the higher expression of T cell-related proinflammatory cytokines such as IL-1β, IL-6, and TNF-can be explained by the lower expression of TGFβ and FoxP3 in TCR Cβ^-/-^ chickens. The higher levels of monocytes in TCR Cβ^-/-^ chickens are associated with inflammation in the spleen and gut, and the infiltration of macrophages into epithelial tissue, leading to the formation of granulomas which occur when the immune system is unable to effectively eliminate persistent antigens (19).

In the TCR Cβ^-/-^ chicken line there is no division between the cortex and medulla of the thymus visible. During maturation, T cells migrate from the cortex into the medulla (25). Whereas αβ T cells need several days to migrate, γδ T cells migrate much faster. This is why Bucy et.al. suggest, that γδ T cells do not undergo the same selection process as αβ T cells (26). Our findings indicate that either the γδ T cells do not need the cortex for the maturation process or the αβ T cells are responsible for the separation of the thymus and therefore the γδ T cells cannot mature. Also, we found that the bursa development is impacted by the knockout because smaller B cell follicles are found in the TCR Cβ^-/-^ chickens. Whether this effect is caused by the loss of CD4^+^ T cells, needs to be further investigated. The avian spleen serves as the primary site for interaction between CD4^+^ T cells and B cells, where after antigenic stimulation, B cells give rise to germinal centers (23). The presence of αβ T cells is necessary for this process, although reports of spontaneous germinal center formation in TCR ^-/-^ mice have been documented. Conversely, in TCRβ^-/-^ mice germinal center formation was absent (21), a phenomenon also observed in the spleens of TCR Cβ^-/-^ chickens in the current study.

The gut-associated lymphatic tissue (GALT) is an important part of the chicken immune system. It has been shown that leukocytes can be regulated by the microbiome of the chicken (27). Herewith it was shown that also the missing αβ T lymphocytes can influence the microbiome, as we found significant differences in the microbiome of TCR Cβ^-/-^ chicken compared to their wild type siblings. In TCR ^-/-^ mice it was shown, that only specific pathogen free housed KO mice develop colitis, but not germ-free mice, concluding that a microbial agent activates the immune system to cause a spontaneous autoimmune reaction (17). This can also explain the inflammation in the stomach of the TCR Cβ^-/-^ chicken.

The results showed that the knockout of γδ T cells does not result in a pronounced phenotype, whereas the knockout of αβ T cells leads to a severe phenotype including granulomas on comb, leg and beak, inflammations of the spleen and the proventriculus, and impaired B cell function and immunoglobulin production due to the loss of CD4^+^ T cells including T_reg_ cells. These findings highlight the crucial role of αβ T cells in regulating the immune response and demonstrate their importance in the chicken immune system. These genetically modified chickens will serve as a tool to study the nature and function of T cell subpopulations in detail by performing various infection experiments. Understanding the distinct functions of γδ and αβ T cells in chickens will help to improve our knowledge of the chicken’s immune system and to develop new strategies for controlling diseases in chickens by targeting specific components of the immune system. Additionally, these genetically modified chicken lines provide a model for understanding the functions of T cells in other species, including humans, which can deepen our understanding of the evolution of the immune system and its role in protecting against pathogens.

## Materials & Methods

### Animals

White Leghorn (Lohmann selected White Leghorn (LSL), Lohmann-Tierzucht GmbH, Cuxhaven, Germany) chickens were used. Animal experiments were approved by the government of Upper Bavaria, Germany (ROB-55.2-2532.Vet_02-17-101 & 55.2-1-54-2532-104-2015). Experiments were performed according to the German Welfare Act and European Union Normative for Care and Use of Experimental Animals. All animals received a commercial standard diet and water *ad libitum*. Genetically modified animals were generated as previously described (12). Shortly, PGCs, with the desired genetic modification, were injected into the vasculature of 65 h old embryos transferred into a turkey surrogate eggshell, and incubated until the hatch of chimeric roosters. Upon sexual maturity, sperm was collected for DNA isolation and genotyping. The germline-positive roosters were bred with wild type hens, first to obtain heterozygous animals, and then siblings were bred together to obtain homozygous animals.

### Genotyping assays

For the TCR Cβ^-/-^ chickens’ primers were designed to detect the TCR Cβ^-/-^ knockout: Forward: 5’ GGTTCGAAATGACCGACCAAGC 3’; Reverse: 5’ GGCTTGCACACTCAGCTCTATAG 3’. A second primer pair was used to detect the TCR Cβ wild type allele: Forward: 5’ GGTTCGAAATGACCGACCAAGC 3’; Reverse: 5’ CACACCATTCACCTTCCAGAC 3’. FIREPol Multiplex DNA Polymerase Mastermix (Solis Biodyne, Tartu, Estonia) was used according to manufactures instructions with an annealing temperature of T_m_ 58°C. For the TCR Cγ^-/-^ chickens’ primers were designed to detect the TCR Cγ^-/-^ knockout: Forward: 5’ GCCATTCCTATTCCCATCCTAAGT 3’; Reverse: 5’ GGTTCGAAATGACCGACCAAGC 3’. A second primer pair was used to detect the wild type constant region of the TCR chain: Forward: 5’ GAGCTCCACGCCATGAAACCATAG 3’; Reverse: 5’ GTTGTCACTGTCACTGGCTG 3’. FIREPol Multiplex DNA Polymerase Mastermix (Solis Biodyne, Tartu, Estonia) was used according to manufactures instructions with an annealing temperature of T_m_ 60°C.

### Flow cytometry

PBMCs were isolated using histopaque density gradient centrifugation (Sigma, Taufkirchen, Germany). 1×10^6^ cells were used per sample and washed with 1% BSA in PBS + 0,01% NaN_3_. Cells were first washed and then stained with primary antibodies (Table S1) for 20 min in the dark on ice. Subsequently, cells were washed and incubated with secondary antibodies (Table S1) for 20 min in the dark on ice.

Thereafter cells were again washed and analyzed using an AttuneNXT flow cytometer (LifeTechnologies, Carlsbad, USA). Data were analyzed with FlowJo 10.8.1 software (FlowJo, Ashland, USA). For the FACS analysis of organs, the organ was strained through a 100µm cell strainer into a falcon holding 5mL PBS. The single cell solution was then further processed as described above.

### ELISA

ELISA was performed as described before (13). Used antibodies are listed in Table S4. OD was measured at 450 nm with FluoStar Omega (Version 5.70 R2 BMG LABTECH, Ortenberg, Germany).

### Histology

Tissue was frozen at -80°C in O.C.T. Tissue Tek Compound (Thermo Fisher Scientific, Waltham, USA). Before sectioning the tissue was stored overnight at -20°C. 7-8µm sections were prepared. For H&E Histology sections were stained with Mayer’s hematoxyline (Medite, Burgdorf, Germany) followed by Eosin (Medite, Burgdorf, Germany). For mounting VectaMount Express Mounting Medium (Biozol, Eching, Germany) was used. Slides were scanned using a Precipoint Microscope. (Version 1.0.0.9628 PreciPoint, Freising, Germany)

For immunohistology, sections were stained with the antibodies listed in Table S2. Antibodies were detected using the Vectastain ABC Peroxidase Kit (Biozol, Eching, Germany) followed by the Vector DAB Kit (Biozol, Eching, Germany). Sections were counterstained with Mayer’s hematoxyline (Medite, Burgdorf, Germany). For mounting VectaMount Express Mounting Medium (Biozol, Eching, Germany) was used. Slides were scanned using Precipoint Microscope. (Version 1.0.0.9628). For fluorescence histology, sections were stained with the following antibodies shown in Table S3. Cell nuclei were stained with 0,01% DAPI (Applichem, Darmstadt, Deutschland). Vectashield Mounting Medium (Biozol, Eching, Germany) was used to mount the stained sections. Images were taken using an ECHO Revolve Microscope. (Revolve Software Version 4.0.5 Discover Echo Inc., San Diego, USA)

### RNA extraction

Bursa, spleen, and thymus were collected 14 days post hatch. Samples were frozen in RNAlater (Sigma, Taufkirchen, Germany) at -20° until further processing. After defrosting samples were rinsed with PBS and homogenized using a SpeedMill Homogenisator (Analytik Jena, Jena, Germany) for 6x 60 seconds. Thereafter the samples were processed according to the Manufacturer’s Protocol for ReliaPrep RNA Tissue Miniprep System (Promega, Fitchburg, USA). PBMCs were isolated using histopaque density gradient centrifugation (Sigma, Taufkirchen, Germany). Further processing was according to the Manufacturer Protocol using ReliaPrep RNA Cell Miniprep System (Promega, Fitchburg USA). RNA integrity was analyzed using an Agilent 2100 Bioanalyzer (Agilent Technologies, Santa Clara, USA). Only RNA with a RIN ≥7,5 was used for downstream analysis.

### cDNA Synthesis

Using Go Script cDNA Kit (Promega, Fitchburg USA) RNA samples were processed according to the manufacturer’s protocol.

### qRT-PCR

Primers (Table S5) were adapted from literature or designed using Benchling (San Francisco, USA). Promega GoTaq qRT-PCR kit (Promega, Fitchburg USA) was used according to the manufacturer’s instructions. Expression was measured using QuantStudio5 (QuantStudio™ Design&analysis Software v1.5.2, Thermo Fisher Scientific, Waltham, USA)

### Morphometry

Sections from duodenum, jejunum, ileum, and caecum were taken from TCR Cγ^-/-^ (n≥3) and wild type chickens (n≥5) 35 days after hatch and frozen in O.C.T. Tissue Tek Compound (Thermo Fisher, Waltham, USA) at -80°C. Before cutting, the tissue was stored overnight at -20°C. 7-8µm sections were prepared. H&E Stain was performed as described above. Measurements were done with the Precipoint Microscope (Version 1.0.0.9628 PreciPoint, Freising, Germany). From each slide, the longest villus and the corresponding crypt were measured and for the Tunica muscularis 4 sites were measured. At least 3 sections per organ and animal were prepared.

### Microbiome analysis

Feces was taken from 14-day-old TCR Cβ^-/-^ chicken and wild type chickens. The feces were collected using a sterile spoon attached to a fecal tube. Feces samples were stored in 1 mL of Stool Stabilizer (Invitek, Berlin, Germany). Caecum content was taken from TCR Cβ^-/-^ animals after 14 days and from TCR Cγ^-/-^ animals after 35 days both compared to wild type. Caecum content was flushed into a tube holding 1 mL Stool Stabilizer (Invitek, Berlin, Germany). From all samples, DNA was extracted and processed as previously described (28).

### Statistical analysis

Statistical analyses were performed using SPSS statistics software (version 28.0.1.1. IBM, Armonk, USA). Normally distributed data (Shapiro-Wilk test p<0.05) were analyzed by student’s T-Test and Mann-Whitney-U Test was applied for not normally distributed data. All P-values <0.05 were marked as significant. Graphs were designed with GraphPad Prism (Version 9.3.1. Dotmatics, Boston, USA). Statistics of the microbiome analysis were performed using Wilcoxon-Rank-Sum Pairwise test.

## Data availability statement

The data that supports the findings of this study is available from the corresponding author upon reasonable request. Raw data for the microbiome analysis are available on SRA (Accession Number: PRJNA934268).

### Ethics approval statement for human and/or animal studies

All animal work was conducted according to relevant national and international guidelines for humane use of animals. Animal experiments were approved by the government of Upper Bavaria, Germany (ROB-55.2-2532.Vet_02-17-101 & 55.2-1-54-2532-104-2015).

### Author contributions

TvH performed and analyzed the experiments and wrote the paper. BS supervised the work and wrote the paper. HS performed experiments and wrote the paper. MA planned experiments and wrote the paper. RK, CZ, DA, AS, KL, H-KV, and TM performed experiments. All authors approved the submitted version.

Acknowledgment

This project was funded by the Deutsche Forschungsgemeinschaft (DFG, German Research Foundation) in the framework of the Research Unit ImmunoChick (FOR5130) project SCHU 2446/6-1 (awarded to BS) and an Emmy-Noether research fellowship (DFG Schu2446/3-1 awarded to BS). We thank Bernd Kaspers, Department of Animal Sciences, Ludwig-Maximilians University Munich for his valuable advises. Also, we thank Silvia Mitterer, Chair of Anatomy, Histology and Embryology, Ludwig-Maximilians University Munich for help with the histology. The anti-GRL1 antibody developed by Thomas, J.-L was obtained from the Developmental Studies Hybridoma Bank, created by the NICHD of the NIH and maintained at The University of Iowa, Department of Biology, Iowa City, IA 52242.

