## supplementary information for "Knockout of αβ but not γδ T cells in chickens is associated with high cytotoxicity and deficiency of regulatory and helper T cells"

**This PDF file includes:**

Figures S1 to S6
Tables S1 to S5
SI References

**Fig. S1:**

A

| Germline transmission | KO with selectable marker cassette | KO w/o selectable marker cassette |
|---|---|---|
| TCR Cγ | 5,3 % | 9 % |
| TCR Cβ | 1,5 % | 5,9 % |

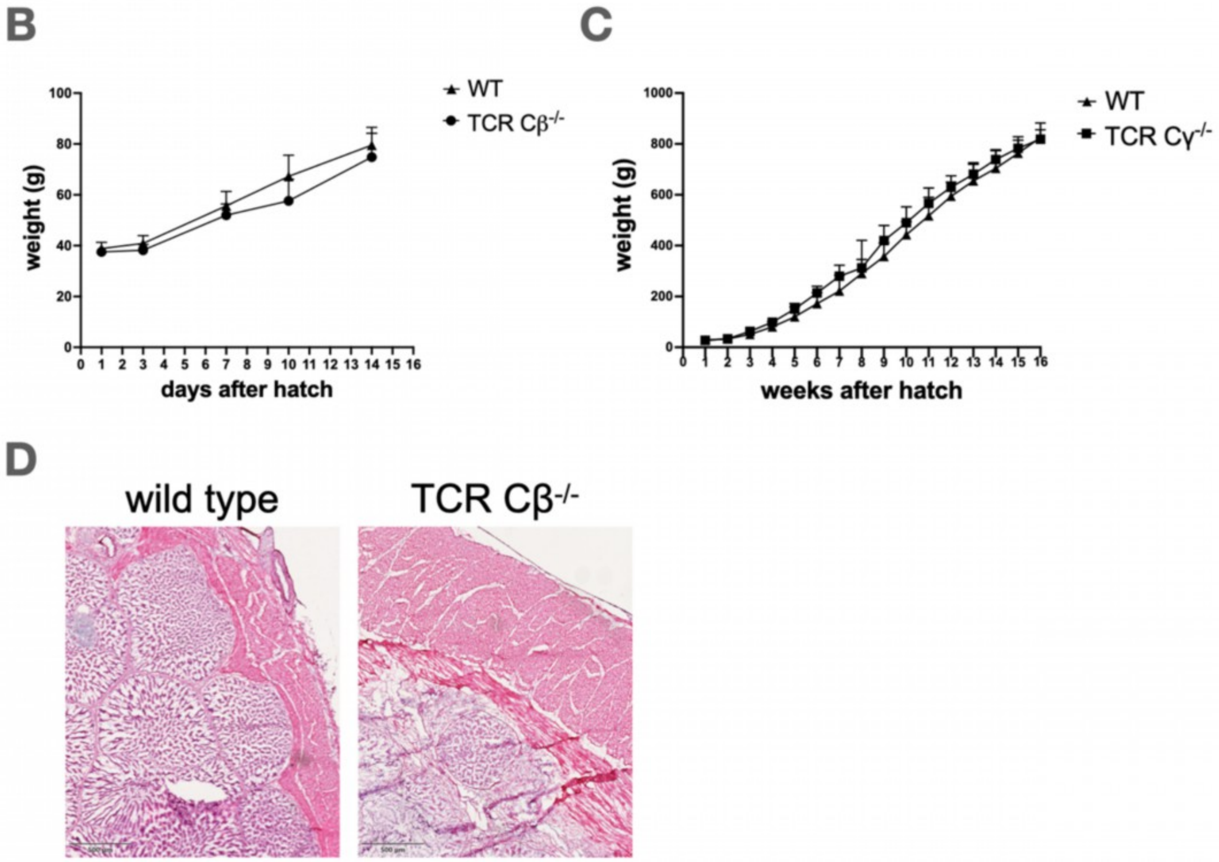


(A) Germline transmission data of TCR C$\gamma$ and TCR C$\beta$ knockout PGCs with and without selectable marker cassette is shown. (B) TCR C$\gamma^{-/-}$ (n=3) and wild type (n=3) animals up to 16 weeks after hatch and (C) TCR C$\beta^{-/-}$ (n≥6) and wild type (n≥5) animals up to 14 days after hatching was monitored. (D) The glandular stomach of a 14-day-old wild type and TCR C$\beta^{-/-}$ chicken were dissected and stained with H&E. One representative picture of at least three animals per genotype is shown. Scale bar = 100µm

**Fig. S2:**

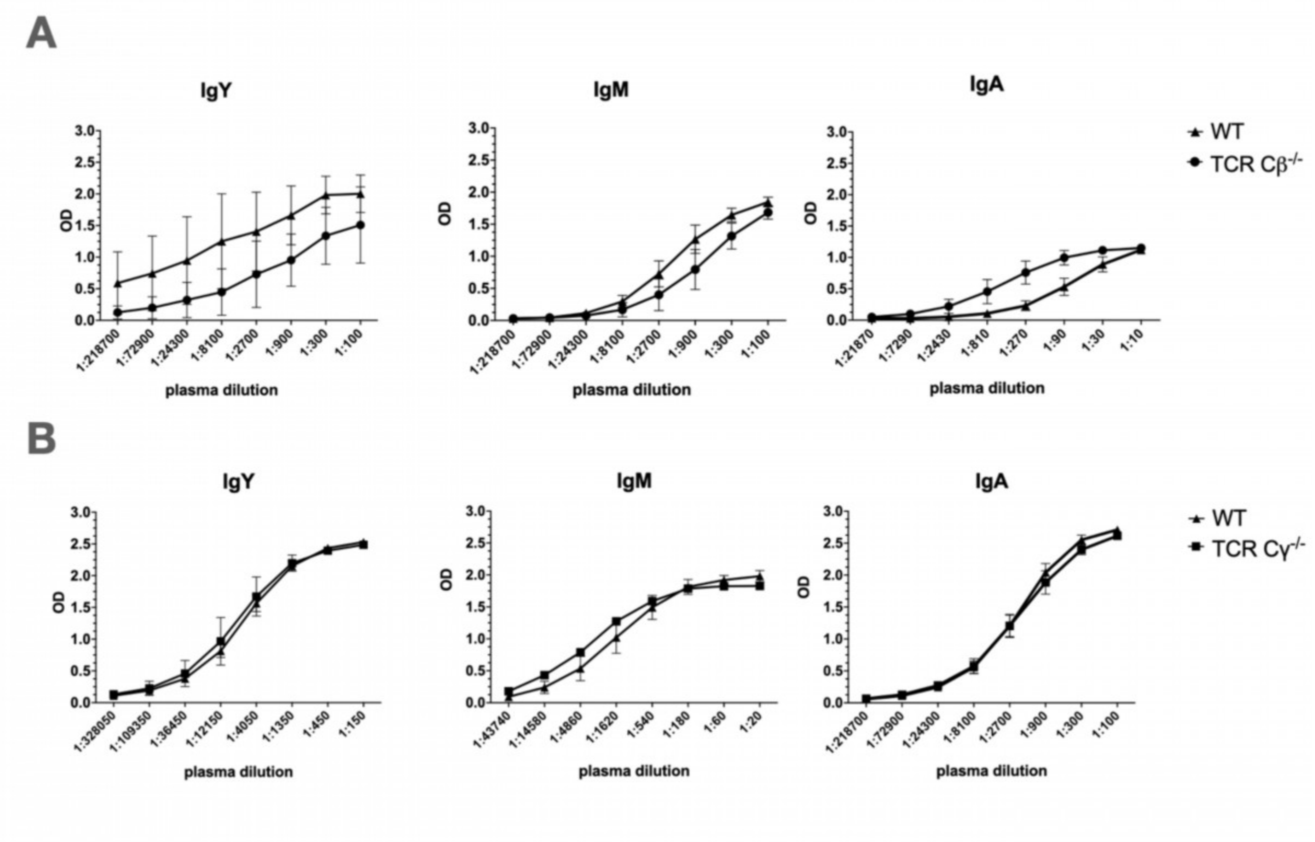


Total plasma IgM, IgY, and IgA immunoglobulin levels of (A) TCR C$\beta^{-/-}$ (n=10) and wild type (n=12) chickens at 14-days- of age and of (B) TCR C$\gamma^{-/-}$ (n=5) and wild type (n=5) chickens at an age of 49 days post-hatch were measured by ELISA. Mean and standard deviation are shown.

**Fig. S3:**

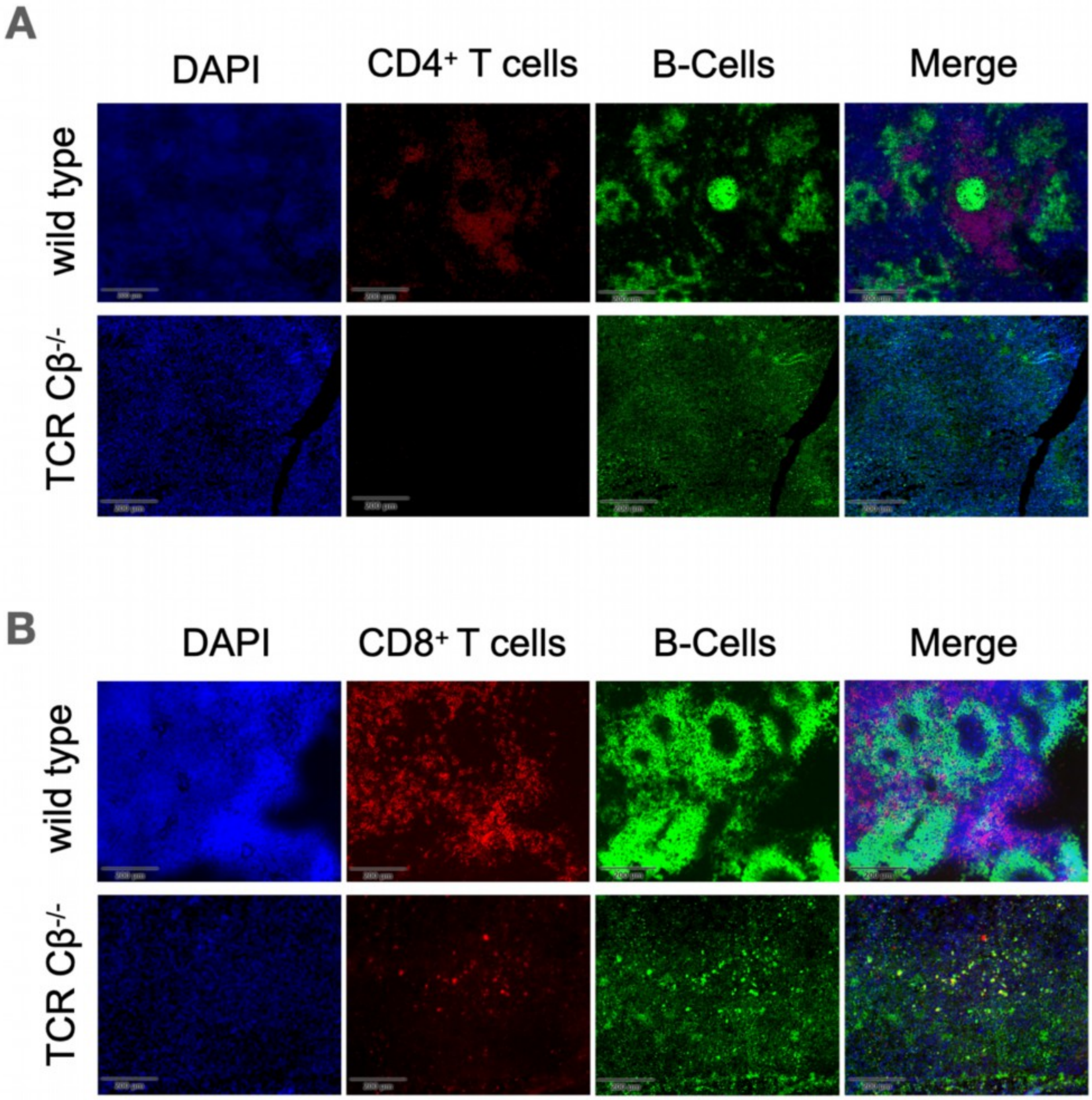


Sections of the spleen of 14-day-old wild type and TCR C$\beta^{-/-}$ chickens were prepared and stained with (A) mouse-anti-chicken-CD4-AF568 (red) and anti-chicken-Bu1-FITC (green) or (B) anti-chicken-CD8-AF568 (red). Nuclei were counterstained with DAPI (blue). One representative image of at least three different animals per genotype is shown. Scale bar = 200μm

**Fig. S4:**

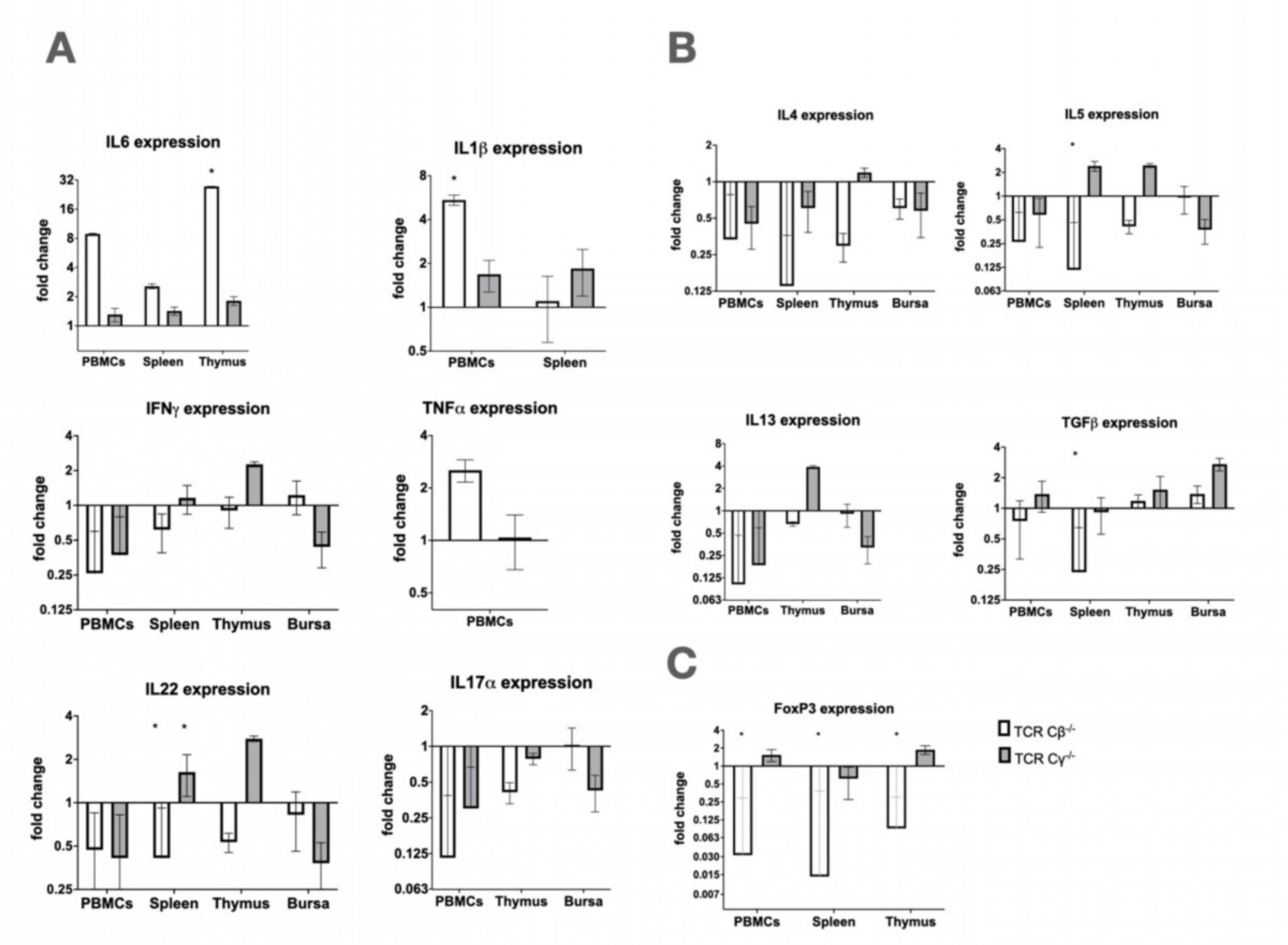


Expression levels of (A) IL-6, IL-1$\beta$, IFN-$\gamma$, TNF$\alpha$, IL-22, IL-17a (B) IL-4, IL-5, IL13, TGF$\beta$ and (C) FoxP3 were analyzed by qRT-PCR. RNA integrity was measured and only RNA with a RIN≥7,5 was used for downstream analysis. qRT-PCR was performed in duplicates. n≥3 animals per genotype at an age of 14 days were used. Mean and standard deviation are shown. * p<0,05

**Fig. S5:**

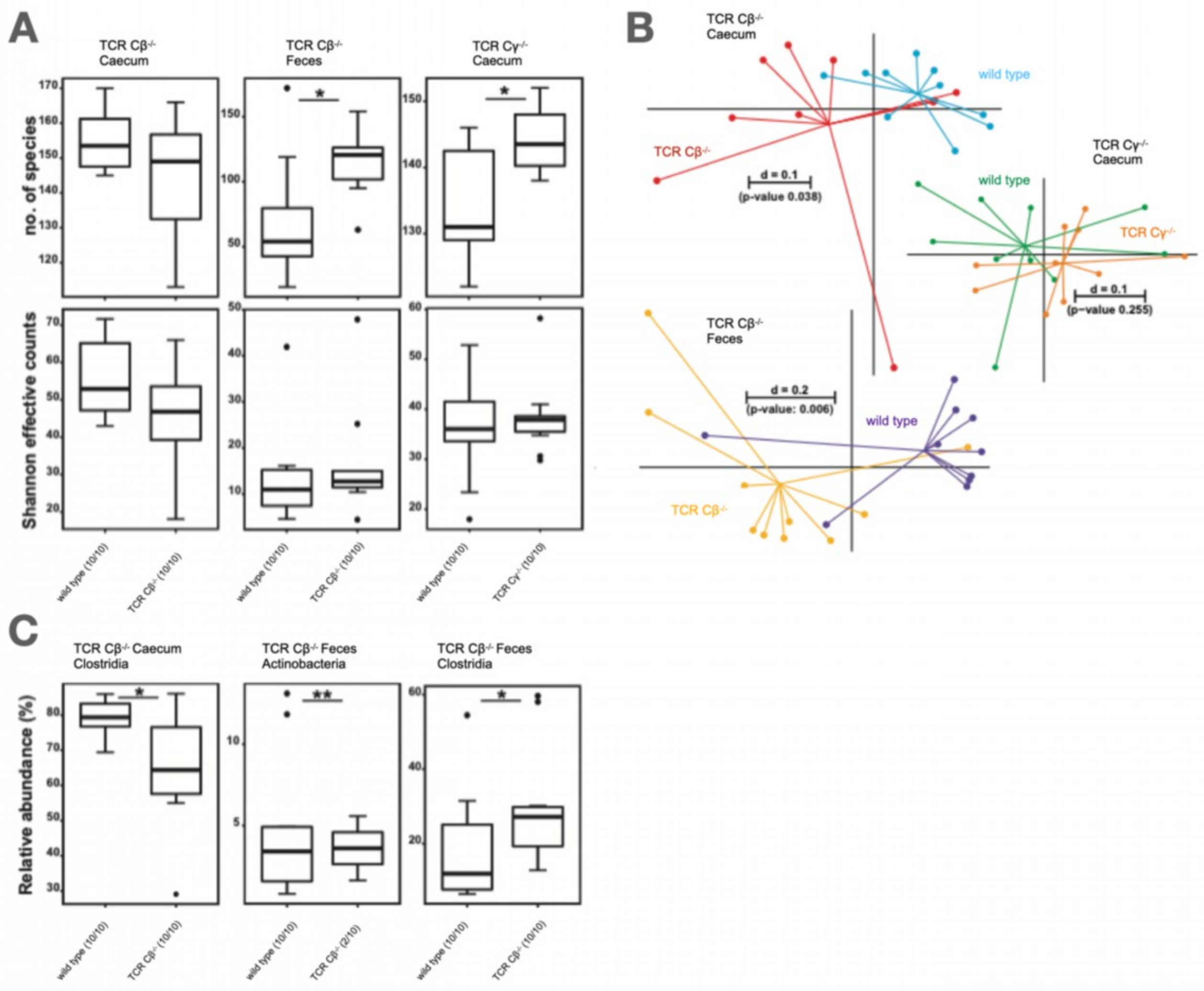


The gut microbiome of TCR $C\beta^{-/-}$ and TCR $C\gamma^{-/-}$ chickens were compared to wild type chickens by 16S rRNA gene amplicon sequencing. Caecum and feces samples were taken 14 days post-hatch from TCR $C\beta^{-/-}$ animals and caecum samples were taken from TCR $C\gamma^{-/-}$ and wild type animals 35 days post-hatch. (A) The number of species and the Shannon effective count are displayed as well as (B) the $\beta$-Diversity and (C) significant differences in relative abundance for Clostridia (class) in caecum and feces of TCR $C\beta^{-/-}$ and for Actinobacteria (class) in feces of TCR $C\beta^{-/-}$. Mean and standard deviation are shown. * $p<0.05$

**Fig. S6:**

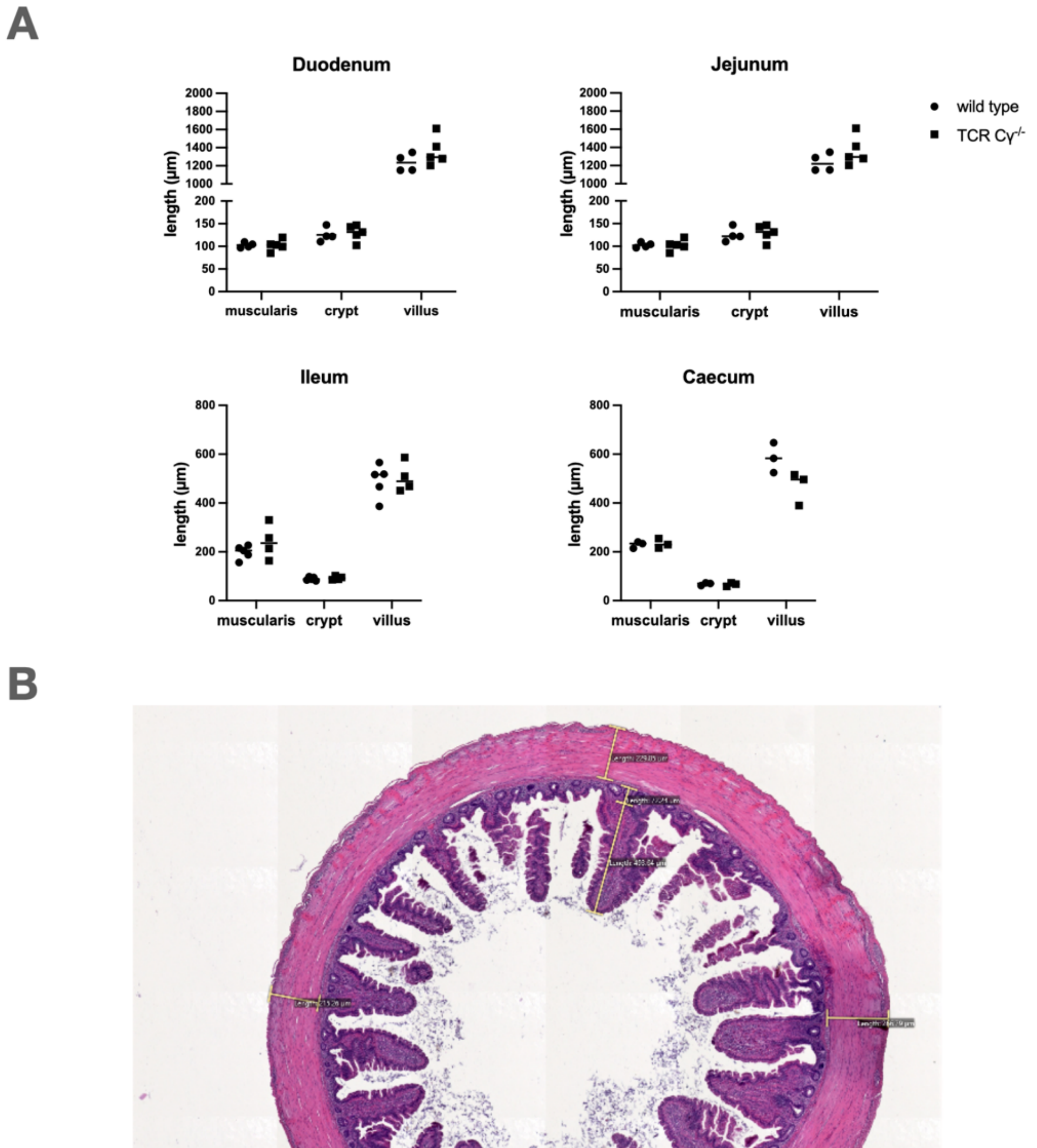


Morphometric analysis of the intestine from TCR Cγ$^{-/-}$ compared to wild type. (A) Length of the tunica muscularis, crypt, and villus from the duodenum, jejunum, ileum, and caecum. (B) Exemplary schema of the caecum, how all intestinal sections were measured. A minimum of 3 intestinal sections were prepared from each animal. n≥3 per genotype at an age of 35 days were used. Mean and standard deviation are shown. * $p<0.05$

**Fig. S7:**

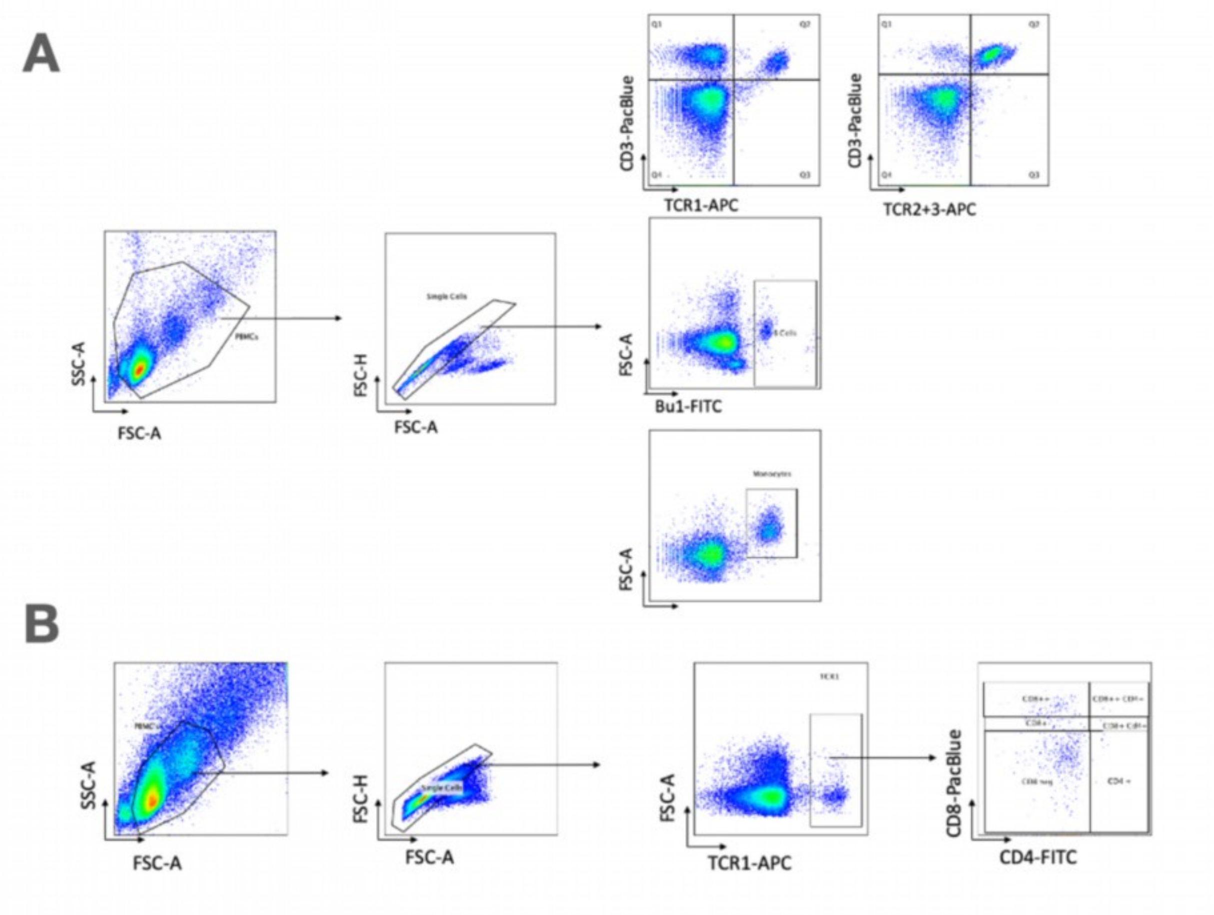


The gating strategy for the flow cytometry analysis is shown (A) for different PBMC populations (CD3, TCR1 / TCR2+3, Bu1, and KUL01) and (B) the TCR1 subpopulations $CD8^{+high}$, $CD8^{+dim}$, $CD^{-neg}$, $CD8^{+high}CD4^{+}$, $CD8^{+dim}CD4^{+}$. One representative example is shown.

**Table S1:** Antibodies used in Flow cytometry

| Antibody name | Manufacturer | Clone | Concentration |
|---|---|---|---|
| mouse anti-chicken CD8a_PacBlue | Biozol | CT-8 | 0,625 µg/mL |
| mouse anti-chicken CD8ß_UNLAB | Biozol | EP42 | 0,625 µg/mL |
| mouse anti-chicken CD4_FITC | Biozol | CT-4 | 0,625 µg/mL |
| mouse anti-chicken TCRgd-BIOT | Biozol | TCR-1 | 0,625 µg/mL |
| mouse anti-chicken TCRab/Vb1-BIOT | Biozol | TCR-2 | 2,5µg/mL |
| mouse anti-chicken TCRab/Vb2-BIOT | Biozol | TCR-3 | 2,5µg/mL |
| mouse anti-chicken Bu1-FITC | Biozol | AV-20 | 2,5µg/mL |
| mouse anti- chicken KUL01 | Biozol | KUL01 | 0,625 µg/mL |
| rat anti- mouse IgG2a_PE | Biozol | SB84a | 0,125µg/mL |
| Streptavidin-APC | VWR | - | 0,2µg/mL |
| goat anti-mouse IgG (H+L)-APC | Biozol | polyclonal | 0,625 µg/mL |

**Table S1:** Antibodies used for Immunohistology

| Antibody name | Manufacturer | clone | Concentration |
|---|---|---|---|
| mouse anti-chicken TCR1 | Biozol | TCR-1 | 2,5µg/mL |
| mouse anti-chicken GRL1 | DSHB | GRL-1 | 0,61µg/mL |
| Mouse anti-chicken Bu1 | Biozol | AV-20 | 5µg/mL |
| Mouse anti-chicken KUL01 | Biozol | KUL01 | 5µg/mL |

**Table S2:** Antibodies used for Fluorescence Histology

| Antibody name | Manufacturer | clone | Concentration |
|---|---|---|---|
| CD4-UNLB | Biozol | CT-4 | 0,01mg/mL |
| CD8a-UNLB | Biozol | CT-8 | 5μg/mL |
| Bu-1-FITC | Biozol | AV-20 | 5μg/mL |
| anti-mouse-IgG-AF568 | Fisher Scientific | polyclonal | 0,01mg/mL |

**Table S3:** Antibodies used for ELISA

| Antibody name | Manufacturer | clone | Concentration |
|---|---|---|---|
| Mouse anti-chicken IgA-UNLB | Biozol | A-1 | 2µg/mL |
| Rabbit anti-chicken IgY | Jackson | polyclonal | 2µg/mL |
| Goat anti-chicken IgM | Biomol | polyclonal | 2µg/mL |
| Goat anti-chicken IgA-HRP | Biomol | polyclonal | 0,1ng/mL |
| Goat anti-chicken IgM-HRP | Biomol | polyclonal | 0,05ng/mL |
| Rabbit anti-chicken IgY-HRP | Jackson | polyclonal | 0,02µg/mL |

**Table S4**: qRT-PCR Primer

| Primer | Sequence 5' to 3' | Tm | Amplicon (bp) | Source | Accession number |
|---|---|---|---|---|---|
| FoxP3 sense | AGTACGCCACAACCTGAGCCT | 59 °C | 157 | Adapted from Burkhardt et al. 2022 (1) | MT133687.1 |
| FoxP3 antisense | TTGGGGTCCTCTCAGCTCCGT | 59 °C | 157 | Adapted from Burkhardt et al. 2022 (1) | MT133687.1 |
| IL13 sense | CTGCCCTTGCTCTCCTCTGT | 59 °C | 123 | Adapted from Liu et al. 2010 (2) | AJ621250.1 |
| IL13 antisense | CCTGCACTCCTCTGTTGAGCTT | 59 °C | 123 | Adapted from Liu et al. 2010 (2) | AJ621250.1 |
| IL17A sense | TTTCTGCACATGGGAAGGTG | 59 °C | 144 | Adapted from Khampeerathuch et al. 2018 (3) | AJ493595 |
| IL17A antisense | CCTGGTTCATGTTGCTGATGC | 59 °C | 144 | Adapted from Khampeerathuch et al. 2018 (3) | AJ493595 |
| IL22 sense | TGTTGTTGCTGTTTCCCTCTTC | 59 °C | 143 | Adapted from Kim et al. 2012 (4) | NM_001199614.1 |
| IL22 antisense | GCCAAGGTGTAGGTGCGATTCC | 59 °C | 143 | Adapted from Yu et al. 2021 (5) | NM_001199614.1 |
| IL4 sense | GTGCCCACGCTGTGCTTAC | 59 °C | 82 | Adapted from Xu et al. 2015 (6) | AJ621249. |

| | | | | | |
|---|---|---|---|---|---|
| IL4 antisense | AGGAAACCTCTCCCTGGATGTC | 59 °C | 82 | Adapted from Xu et al. 2015 (6) | AJ621249. |
| IL5 sense | GGAACGGCACTGTTGAAAAATAA | 59 °C | 111 | Adapted from Liu et al. 2010 (2) | NM_001007084.2 |
| IL5 antisense | TTCTCCCTCTCCTGTCAGTTGTG | 59 °C | 111 | Adapted from Liu et al. 2010 (2) | NM_001007084.2 |
| IL6 sense | GCTTCGACGAGGAGAAATGC | 59 °C | 139 | Breithaupt 2011 (7) | NM_204628 |
| IL6 antisense | GCCAGGTGCTTTGTGCTGTA | 59 °C | 139 | Breithaupt 2011 (7) | NM_204628 |
| TGFβ sense | CGGCCGACGATGAGTGGCTC | 59 °C | 120 | Adapted from Brisbin et al. 2010 (8) | M31160.1 |
| TGFβ antisense | CGGGGCCCATCTCACAGGGA | 59 °C | 120 | Adapted from Brisbin et al. 2010 (8) | M31160.1 |
| TNFα sense | TGCTGTTCTATGACCGCC | 59 °C | 174 | Adapted from Farag et al. 2021 (9) | NM_204267.2 |
| TNFα antisense | CTTTCAGAGCATCAACGCA | 59 °C | 174 | Adapted from Farag et al. 2021 (9) | NM_204267.2 |
| IFNγ sense | CACTGACAAGTCAAAGCCGCAC | 59 °C | 129 | designed using Benchling | NM_205149.2 |
| IFNγ antisense | AAGTCGTTCATCGGGAGCTTGG | 59 °C | 129 | designed using Benchling | NM_205149.2 |
| IL1β sense | GTGAGGCTCAACATTGCGCTGTA | 59 °C | 214 | Adapted from Brisbin et al. 2010 (8) | NM_204524.2 |

| | | | | | |
|---|---|---|---|---|---|
| IL1β antisense | TGTCCAGGCGGTAGAAGATGAAG | 59 °C | 214 | Adapted from Brisbin et al. 2010 (8) | NM_204524.2 |
| r18S sense | CATGTCTAAGTACACACGGGCGGTA | 59 °C | 136 | Laparidou et.al 2019 (10) | NC_052547.1 |
| r18S antisense | GGCGCTCGTCGGCATGTATTA | 59 °C | 136 | Laparidou et.al 2019 (10) | NC_052547.1 |
